# THESEUS1 modulates cell wall stiffness and abscisic acid production in *Arabidopsis thaliana*

**DOI:** 10.1101/2021.07.22.453418

**Authors:** Laura Bacete, Julia Schulz, Timo Engelsdorf, Zdenka Bartosova, Lauri Vaahtera, Guqi Yan, Joachim Gerhold, Tereza Tichá, Camilla Øvstebø, Nora Gigli-Bisceglia, Svanhild Johannessen-Starheim, Jeremie Margueritat, Hannes Kollist, Thomas Dehoux, Scott A.M. McAdam, Thorsten Hamann

**Author notes:** Corresponding author: Mail. Laboratory of Plant Physiology, Wageningen University, Radix Building, 6708 PB Wageningen, The Netherlands. Research center for non-destructive testing GmbH, Science Park 2, Altenberger Strasse 69, A 4040 Linz, Austria (Guqi Yan, Nora Gigli-Bisceglia).

## Abstract

Plant cells can be distinguished from animal cells by their cell walls and high turgor pressure. Although changes in turgor and stiffness of cell walls seem coordinated, we know little about the mechanism responsible for coordination. Evidence has accumulated that plants, like yeast, have a dedicated cell wall integrity maintenance mechanism. This mechanism monitors the functional integrity of the wall and maintains it through adaptive responses when cell wall damage occurs during growth, development, and interactions with the environment. The adaptive responses include osmo-sensitive-induction of phytohormone production, defence responses as well as changes in cell wall composition and structure. Here, we investigate how the cell wall integrity maintenance mechanism coordinates changes in cell wall stiffness and turgor in *Arabidopsis thaliana*. We show that the production of abscisic acid (ABA), the phytohormone modulating turgor pressure and responses to drought, depends on the presence of a functional cell wall. We find that the cell wall integrity sensor THESEUS1 modulates mechanical properties of walls, turgor loss point and ABA biosynthesis. We identify RECEPTOR-LIKE PROTEIN 12 as a new component of cell wall integrity maintenance controlling cell wall damage-induced jasmonic acid production. Based on the results we propose that THE1 is responsible for coordinating changes in turgor pressure and cell wall stiffness.

**Significance statement:** Plants need to constantly adapt to a changing environment. This includes responses to biotic and abiotic stress. Key elements influencing the response to abiotic stress are the plant cell walls surrounding all cells and the phytohormone abscisic acid, which influences turgor pressure in plants. Turgor pressure in plant cells is much higher than in animal cells and a key driver of plant growth and development. Here we investigate the mechanism regulating cell wall stiffness and coordinating changes in stiffness and turgor. We characterize key elements of the mechanism and dissect its mode of action. This knowledge will enable us to pursue novel approaches to improve plant resistance to drought stress, which is crucial in a rapidly changing environment.

## Introduction

Plant cell walls are chemically complex structures, changing their composition and structure in a highly dynamic, adaptive manner to meet different biological functions (1). The walls provide structural support during development, are frequently the first line of defense in response to environmental stress and constrain the high turgor pressure in cells, pressures that commonly exceed the levels found in bicycle tires (2–4). Relative abundance and organization of wall components determines the mechanical characteristics of cell walls and varies between plant species, cell types and developmental stages. Major components include cellulose microfibrils (the main load-bearing elements), hemicelluloses, pectins, lignin and cell wall proteins (5). While the mechanical characteristics of epidermal cell walls are well-described, knowledge regarding the mechanical properties of cell walls in sub-epidermal tissues in vivo is limited (3). Recently Brillouin microscopy, which is well-established in materials sciences, has been adapted for use in life sciences to address this lack of knowledge (6, 7). This label free, quantitative microscopy technology is taking advantage of the Brillouin light scattering (BLS) effect. It is based on characterizing the interactions of a laser light with the sample, which involve picosecond-timescale density fluctuations due to spontaneous molecular motions. Since the interactions couple photons to longitudinal phonons in the sample, variations in the scattering spectra of the laser light can be interpreted as response of the sample to an infinitesimal uniaxial compression. This response can be integrated with the refractive index to extract the Brillouin elastic contrast (vB), which reveals changes in stiffness of the sample examined.

Knowledge regarding the mechanisms regulating plant cell wall composition and structure has increased recently. However, our understanding of how changes in composition and structure result in modifications of cell wall stiffness is still limited (3). More importantly we know even less about the mechanisms responsible for controlled changes in cell wall stiffness. The plant cell wall integrity (CWI) maintenance mechanism is of interest in this context because it could contribute to controlling cell wall stiffness in plants. The mechanism monitors the functional integrity of cell walls during growth, development and interactions of plants with their environment (8). Upon impairment of integrity the mechanism initiates adaptive changes in cell wall composition and structure as well as cellular metabolism including induction of jasmonic acid (JA) and salicylic acid (SA) production. Importantly, all these responses can be suppressed by performing co-treatment with osmotica, which reduces turgor pressure levels and is similar to observations made in *Saccharomyces cerevisiae* (9–14). JA has also been implicated in regulating responses to mechano-stimulation in addition to its more established role in biotic stress responses, illustrating the relevance of mechano-perception for biotic stress responses (15). Different components involved in mechano- and turgor-sensing are required for CWI maintenance in *Arabidopsis thaliana* (12). Amongst them are members of the *Catharanthus roseus* receptor-like kinase 1-like (*Cr*RLK1L) family, which contribute to CWI maintenance in different biological processes (16). Two family members, FERONIA (FER) and THESEUS1 (THE1), are of particular interest. THE1 is required for hypocotyl cell elongation, responses to cell wall damage (CWD) induced by inhibition of cellulose biosynthesis and during plant-pathogen interactions (12, 17, 18). THE1 interacts with the peptide ligand RAPID ALAKALINIZATION FACTOR 34 (RALF34) to control also lateral root initiation and is dependent on FER activity (19). FER integrates and coordinates a large number of cellular processes during development and response to stress. The processes include CWI maintenance, coordination of changes in cell wall composition with vacuolar expansion and modulation of Abscisic acid (ABA) production, a phytohormone required for adaptation to drought stress (20–23). FER delivers these highly specific, context-dependent functions by acting as a scaffold protein interacting with a large number of signalling peptides and co-receptors often in a pH-dependent manner (24).

ABA is essential for plant adaptation to drought stress and has therefore been investigated extensively in guard cells (25). ABA biosynthesis is well-characterized and requires enzymes like *ABA DEFICIENT 2* (*ABA2*) (26). Newly produced ABA binds to the PYRABACTIN RESISTANCE 1-LIKE 9 (PYR)-receptor, with the binding inducing a modification, allowing interaction of PYR with ABA INSENSITIVE1 (ABI1) and formation of a complex. The complex activates the kinase OPEN STOMATA1 (OST1), which in turn activates the SLOW ANION CHANNEL1 (SLAC1) through phosphorylation, leading to release of anions (Cl^-^ and NO_3_^-^) causing membrane-depolarization. Depolarization causes release of K^+^-ions through K^+^- channels, resulting in water leaving the guard cell and turgor pressure being reduced (27). This simplified overview highlights the extensive knowledge available about ABA biosynthesis and signaling. However, understanding of the osmo-sensitive processes responsible for induction of ABA production in response to drought is very limited (28, 29).

We hypothesized previously that interactions between *Cr*RLK1Ls and ABA-mediated processes may coordinate adaptive changes in turgor pressure and cell wall stiffness in response to cell expansion or shrinking (8). Here we investigated changes in cell wall stiffness in vivo in subepidermal tissues in Arabidopsis seedling roots, established the function of THE1 in regulation of stiffness and proceeded to show that THE1 also modulates ABA production induced by interactions between the plasma membrane and the cell wall. Using a transcriptomics-based approach, we identified RECEPTOR-LIKE PROTEIN 12 as new component of the CWI maintenance mechanism and characterized itś function in cell wall metabolism, response to CWD and ABA-mediated processes.

## Results

### THE1 modulates cell wall stiffness in Arabidopsis seedling roots

To investigate the interplay between cell wall stiffness and turgor pressure *in vivo*, we treated *A. thaliana* seedlings with the cellulose biosynthesis inhibitor isoxaben (ISX) to cause CWD in a tightly controlled manner and impair CWI, sorbitol to reduce turgor pressure, or a combination of both and studied the effects using Brillouin microscopy (12, 30). Wildtype (Col-0) seedlings were treated with ISX for 6 hours in a pilot study. Longitudinal scans with a Brillouin microscope along the treated seedling roots in three different depths detected the highest BLS contrast in the stele 450 µm from the root tip (SI Appendix, Fig. S1A, B). Subsequently seedling roots were mock-, ISX-, sorbitol-, or ISX/sorbitol-treated for six hours and radial scans were performed in this area. Light microscopy-based studies revealed that the different treatments have pronounced effects on seedling root morphology in this area (SI Appendix, Fig. S1C). Mock and sorbitol-treated roots appeared normal while ISX-treated roots appeared swollen and sorbitol/ISX-treated ones exhibited reduced swelling. Radial scans with a Brillouin microscope revealed in roots treated with ISX or sorbitol pronounced reductions in stiffness in the stele in this area (**Fig. 1**). By contrast, stiffness in Col-0 roots co-treated with ISX and sorbitol was similar to mock-treated controls. In *isoxaben resistant1-1* (*ixr1-1*) mock-, ISX and sorbitol-treated roots stiffness levels differed from wildtype, but responses to treatments were qualitatively equivalent to those observed in Col-0. The observed changes in stiffness might be modulated by dynamic interactions between the plasma membrane and the cell wall. We hypothesized that THE1 could contribute to these interactions since it is membrane-localized and required for responses to ISX (12, 17). Mock-treated *the1-1* (loss of function allele) roots exhibited pronouncedly lower stiffness compared to wildtype, and treatments induced no significant changes compared to mock-treated roots. ISX and sorbitol-treatments of *the1-4* (gain of function allele) roots resulted in pronounced differences compared to mock controls and similar to effects observed in Col-0 roots (31). Importantly, combined ISX/sorbitol treatments resulted in stiffness levels similar to those observed in sorbitol-only treated roots. ISX (cell wall weakening) and sorbitol (turgor reduction) treatments of Col-0 led to similarly reduced cell wall stiffness in sub-epidermal tissue layers, while combined treatments led to enhanced stiffness. These observations suggest that changes in turgor pressure and ISX-treatment affect stiffness through separate processes and that these processes may jointly modulate stiffness in sub-epidermal tissue layers. THE1 seems to be required for regulation of stiffness during growth and following CWD induced by ISX, based on the phenotypes observed in *the1-1* seedling roots. While loss of THE1 has no effect on stiffness changes induced by ISX/sorbitol co-treatment, overactive THE1 prevents these stiffness increases. This suggests that THE1 influences indirectly the process controlling the response to ISX/sorbitol co-treatments, possibly as a consequence of THE1 function in modulating responses to ISX.

**Fig. 1:**
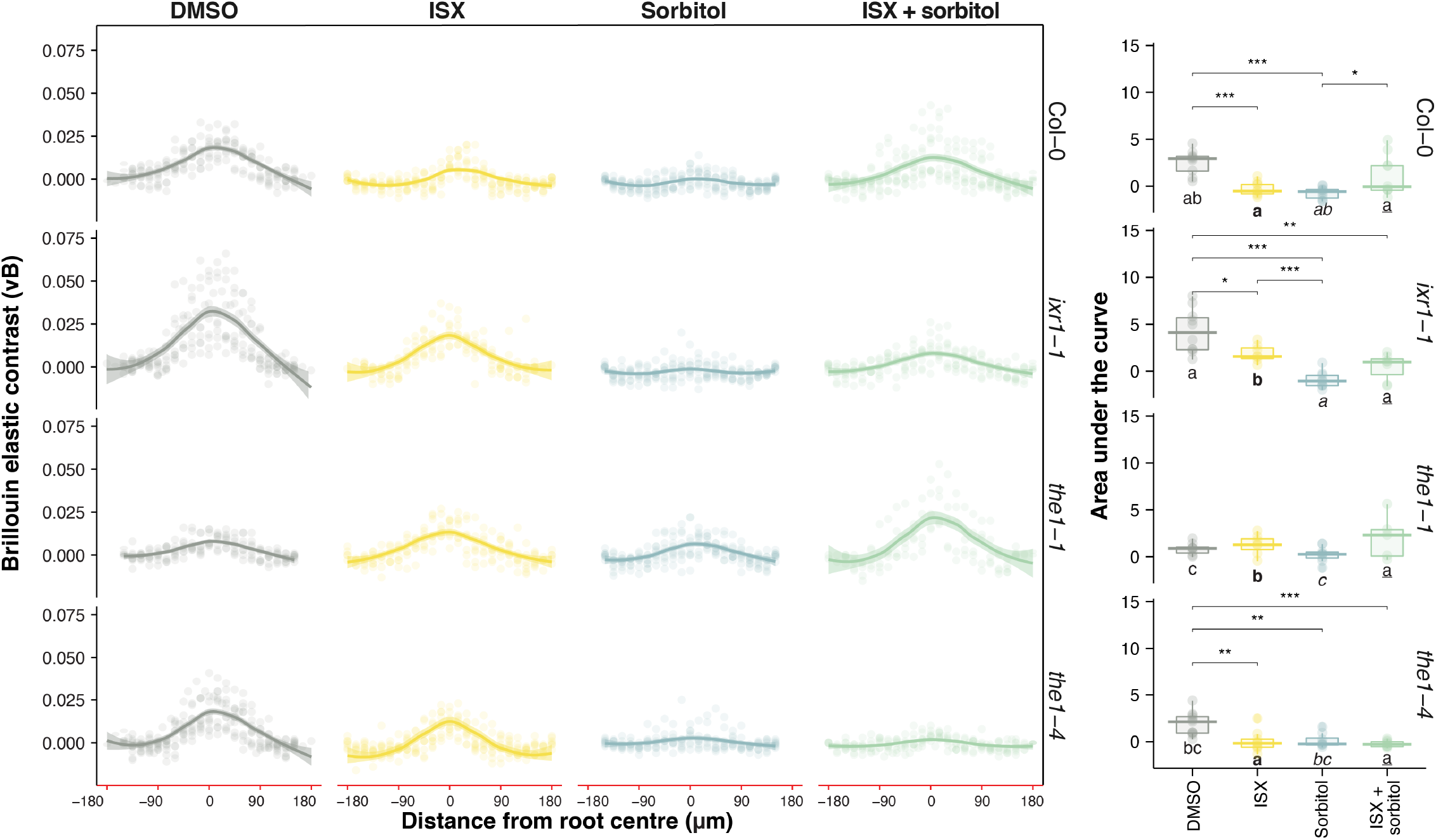
Changes in stiffness of *A. thaliana* root cell walls are mediated by THE1. Brillouin elastic contrast measurements (νB) across the radial axis in the basal meristem–transition zone of *A. thaliana* seedling roots from Columbia-0 (Col-0), *isoxaben resistant1-1 (ixr1-1*), *theseus1-1 (the1-1*) and *the1-4.* Brillouin measurements were performed after 6 hours of treatment as indicated. Dots represent individual measurements (*N* > 10), with lines representing corresponding regression curves (LOESS adjustment) with a 95% confidence interval (shadowed). Areas under the curves were calculated for each root (*N* > 10). Asterisks indicate significance levels allowing comparison of treatment effects within the same genotype (Kruskal–Wallis test with Dunn *post-hoc* analysis, * *P* < 0.05, ** *P* < 0.01, *** *P* < 0.001) whereas letters indicate statistically significant different groups (*P* < 0.05) for the same treatment between different genotypes (i.e., by columns).

### THE1 is a negative regulator of ABA biosynthesis in seedlings

Hyper-osmotic stress induces ABA biosynthesis and reduced turgor pressure, while ISX-treatment triggers jasmonic acid (JA) production in a THE1-dependent manner (9, 32–34). We treated seedling roots as before and used promoter reporter constructs to determine, if JA, ABA levels and *THE1* expression change in the root area where stiffness changes were observed (35–37). Expression of the promoter *pTHE1::YFP* reporter was detectable in mock-treated seedling root tips and induced by ISX-treatment in the region where changes in stiffness had been detected (**Fig. 2A**). *pTHE1::YFP* expression was particularly pronounced after ISX-treatment in the stele and less in the surrounding tissue layers. Sorbitol co-treatment seemed to reduce ISX-induced expression to a level observed in the sorbitol-treated roots. The JA reporter (*pJASMONATE-ZIM-DOMAIN PROTEIN 10* (*pJAZ10)::YFP*) exhibited ISX-induced, sorbitol-sensitive expression in and around the stele of seedling roots in a similar region like *pTHE1*::YFP (**Fig. 2B**). The ABA reporter *pRAB GTPASE HOMOLOG 18-1* (p*RAB18)::GUS-GFP* was preferentially expressed in sub-epidermal cell layers in ISX-, sorbitol- and ISX/sorbitol-treated seedling roots (**Fig. 2C**).

**Fig. 2:**
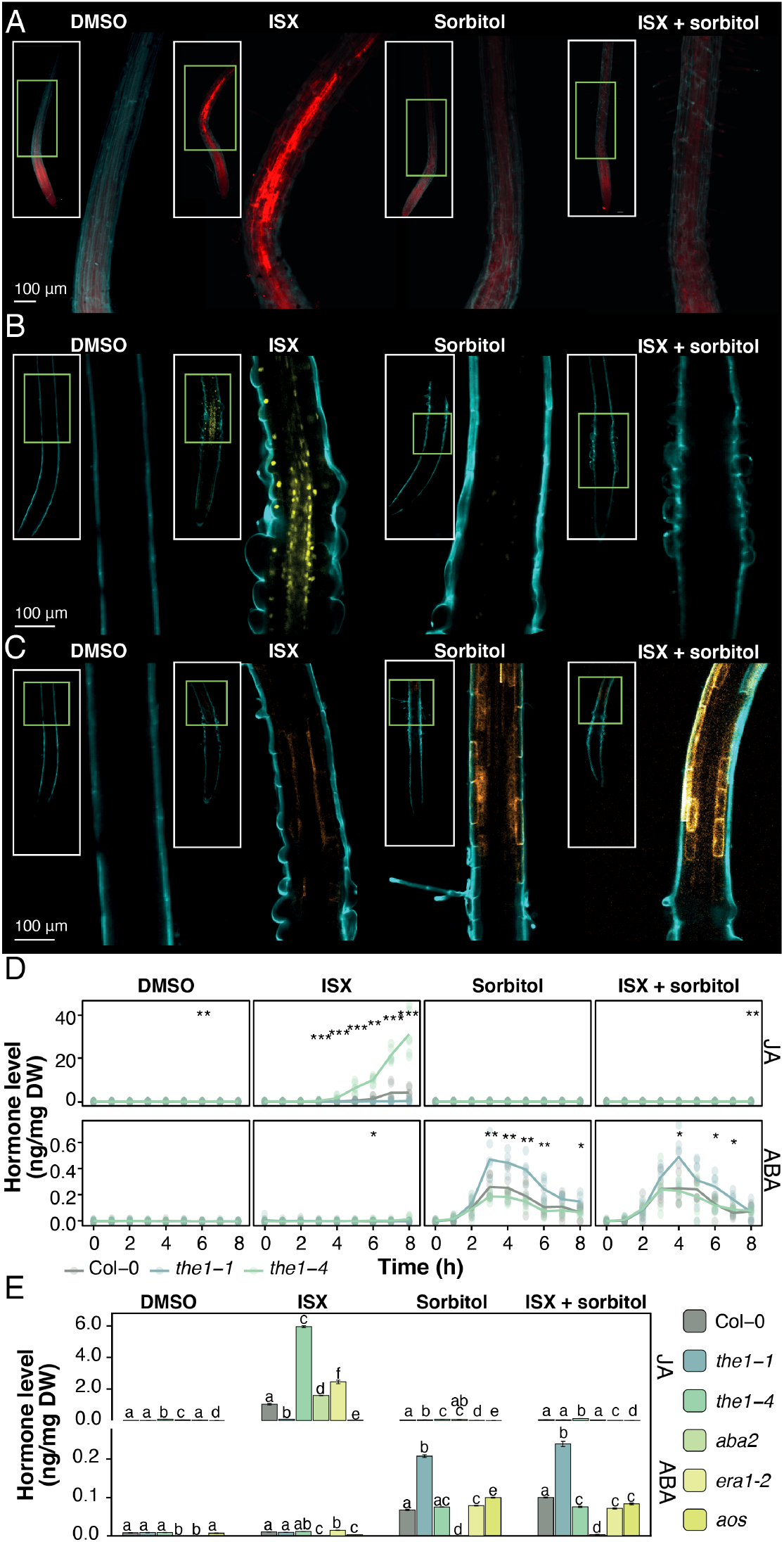
Hyperosmotic and CWI signalling modulate JA / ABA induction and reporter gene expression. (*A-C*) Confocal light microscope images of seedlings expressing *pTHE1*::YFP (red) (*A*), *pJAZ10::YFP* (JA reporter, yellow) (*B*) or *pRAB18::GFP-GUS* (ABA reporter, orange) (*C*) exposed to different treatments as indicated in figure. Root cell walls were counter-stained with Calcofluor-White Solution (cyan). (*D*) Quantification of ABA and JA levels in nanogram hormone per milligram dry weight (DW) Genotypes and treatments indicated in figure. Points represent individual measurements, and lines show average values (*N* = 3). Asterisks indicate statistically significant differences (* *P* < 0.05, ** *P* < 0.01, *** *P* < 0.001, Kruskal–Wallis test). (*E*) Phytohormone levels in whole seedlings with treatments (6 hours) and genotypes indicated in figure; Average values (*N* = 4) ± SEM. Letters indicate significant within-group differences between genotypes (Kruskal–Wallis test with Dunn *post-hoc* analysis, *P* < 0.05).

We proceeded to investigate if THE1 modulates sorbitol- and ISX-induced ABA and JA biosynthesis by performing time course experiments and measuring both phytohormones simultaneously in Col-0, *the1-1* and *the1-4* seedlings (**Fig. 2D**). In mock, sorbitol and ISX/sorbitol treated seedlings JA production was not increased. In ISX-treated *the1-1* seedlings JA production was reduced compared to Col-0 controls while it was enhanced in *the1-4*. ABA production was not increased in mock- or ISX-treated seedlings. In sorbitol and ISX/sorbitol-treated Col-0 and *the1-4* seedlings ABA production was transiently increased in a similar manner. However, in *the1-1* seedlings the increases were enhanced in a pronounced manner. These results suggested that THE1 acts as a positive regulator of JA production in response to CWD and negative regulator of ABA production induced by hyper-osmotic stress.

JA- and ABA-based processes interact to mediate plant adaptation to drought (38). Therefore, we investigated if ISX- or sorbitol-induced JA and ABA production modulate each other and characterized the function of THE1 in this context. We quantified JA and ABA production in *the1-1*, *the1-4*, *aba2*, *enhanced response to aba 1* (*era1*) or *allene oxide synthase* (*aos*, deficient in JA production) seedlings mock-, ISX-, sorbitol-, or ISX/sorbitol-treated for six hours (**Fig. 2E**). In mock-treated seedlings no pronounced changes in JA and ABA production were detected. The effects of manipulating THE1 activity on JA and ABA production were comparable to the results presented in fig. 2D. In ISX-treated *aos* seedlings JA production was not detectable. While ISX-treated *aba2* and *era1* seedlings exhibited slightly enhanced JA production compared to Col-0, the effects were less pronounced than in *the1-4* seedlings. ABA was not detectable in sorbitol and ISX/sorbitol treated *aba2* seedlings. Changes in ABA production were limited in *aos* and *era1-2* after sorbitol or ISX/sorbitol treatments. These results indicate that genetically manipulating ABA and JA biosynthesis affects induction of the corresponding phytohormone by sorbitol or ISX. However, the effects are less pronounced than those observed when manipulating THE1 activity, suggesting that the effects observed are not direct. Interactions between JA and ABA mediated processes responsible for adaptation to drought stress have been described before and could be responsible for the effects we observe here (38).

We followed up further by investigating how manipulation of CWI signalling and ABA biosynthesis affects downstream transcriptional responses induced by ISX or sorbitol. Expression levels of *RAB18* and the ISX-induced *TOUCH4* (*TCH4*) gene were used as readouts (39). We characterized the expression of *RAB18* and *TCH4* in *the1-1*, *the1-4*, and ABA-related (*aba2* and *era1*) mutant seedlings mock-, ISX-, or sorbitol-treated for six hours (SI Appendix Fig. S2). *RAB18* expression did not change in a pronounced way in mock-treated seedlings, was not detectable in *aba2* while a limited expression increase was detectable in ISX-treated *era1-2* seedlings. *RAB18* expression was elevated in sorbitol-treated *the1-1* seedlings but similar to Col-0 in ISX/sorbitol treated *the1-1* seedlings. In *the1-4* seedlings *RAB18* expression was reduced in response to both treatments compared to Col-0 seedlings. In sorbitol-treated *era1-2* seedlings expression was similar to Col-0 and reduced in response to ISX-/sorbitol-treatment. The transcriptional effects observed for *TCH4* were pronouncedly different. In mock-treated seedlings *TCH4* expression was elevated only in *the1-1*. In sorbitol and ISX/sorbitol-treated seedlings all genotypes examined exhibited reduced *TCH4* expression compared to Col-0. Expression of *TCH4* was reduced similarly in *the1-1* and *the1-4* upon ISX-treatment, while it was enhanced both in *aba2* and *era1-2* seedlings. The results indicate that ABA production (*aba2*) is required for *RAB18* expression while ERA1 activity seems not essential. Manipulating THE1 activity has similar effects on *RAB18* expression like on ABA production, reinforcing the notion that THE1 is a negative regulator of ABA-mediated processes. These results imply that *ABA2* and *ERA1* act as negative regulators of *TCH4* expression, while THE1 has apparently no direct regulatory role.

### RLP12 regulates CWD-induced JA production

We performed an RNA-seq experiment to identify additional genes required for CWI maintenance. After growing Col-0 seedlings for six days they were mock-, ISX-, sorbitol- or ISX/sorbitol-treated for one hour. We then proceeded to identify genes whose expression changes in response to ISX- or sorbitol-treatments but does not significantly change in co-treated seedlings (**Fig. 3A**). Expression patterns of 10 genes seemed to meet these criteria with two of them (*PROPEPTIDE1* and *2*) previously implicated in CWI maintenance (12). This group also included *RECEPTOR-LIKE PROTEIN12* (*RLP12*), which belongs to a gene family with 57 members and has been implicated in biotic and osmotic stress responses (40). Time course experiments with mock-, ISX-, sorbitol-, and ISX/sorbitol-treated Col-0 and isoxaben-resistant *ixr1-1* seedlings were performed to validate transcriptomics-derived expression data for *RLP12* using qRT-PCR. ISX induced *RLP12* expression in a sorbitol-sensitive manner in Col-0 but not in *ixr1-1* seedlings (**Fig. 3B**). A *pRLP12::GUS-GFP* reporter construct was generated and seedlings expressing the construct were treated in the same manner for six hours (**Fig. 3C**). Analysis of reporter construct expression suggested that *RLP12* expression is induced in the stele of the root transition/elongation zone (similar to p*THE1::YFP*) and an area in the differentiated seedling root by ISX in a sorbitol-sensitive manner (**Fig. 3C**). In seedlings homozygous mutant for three different T-DNA insertions in *RLP12* ABA and JA production were determined after mock-, ISX-, sorbitol- or ISX/sorbitol-treatments (SI Appendix Fig. S3A, B). Reduced JA production was detected after ISX treatment while ABA production did not exhibit pronounced differences compared to Col-0 controls. In addition, time course experiments were performed to investigate how loss of *RLP12* affects temporal dynamics of phytohormone induction by the different treatments (**Fig. 3D**). JA production was less induced by ISX in *rlp12-1* seedlings than in Col-0 while ABA production did not differ in a pronounced manner. These results indicate that *RLP12* is required for ISX-induced JA production, is expressed in the same region of the seedling root in an ISX-induced, sorbitol sensitive manner like *THE1* and implicate *RLP12* in CWI maintenance.

**Fig. 3:**
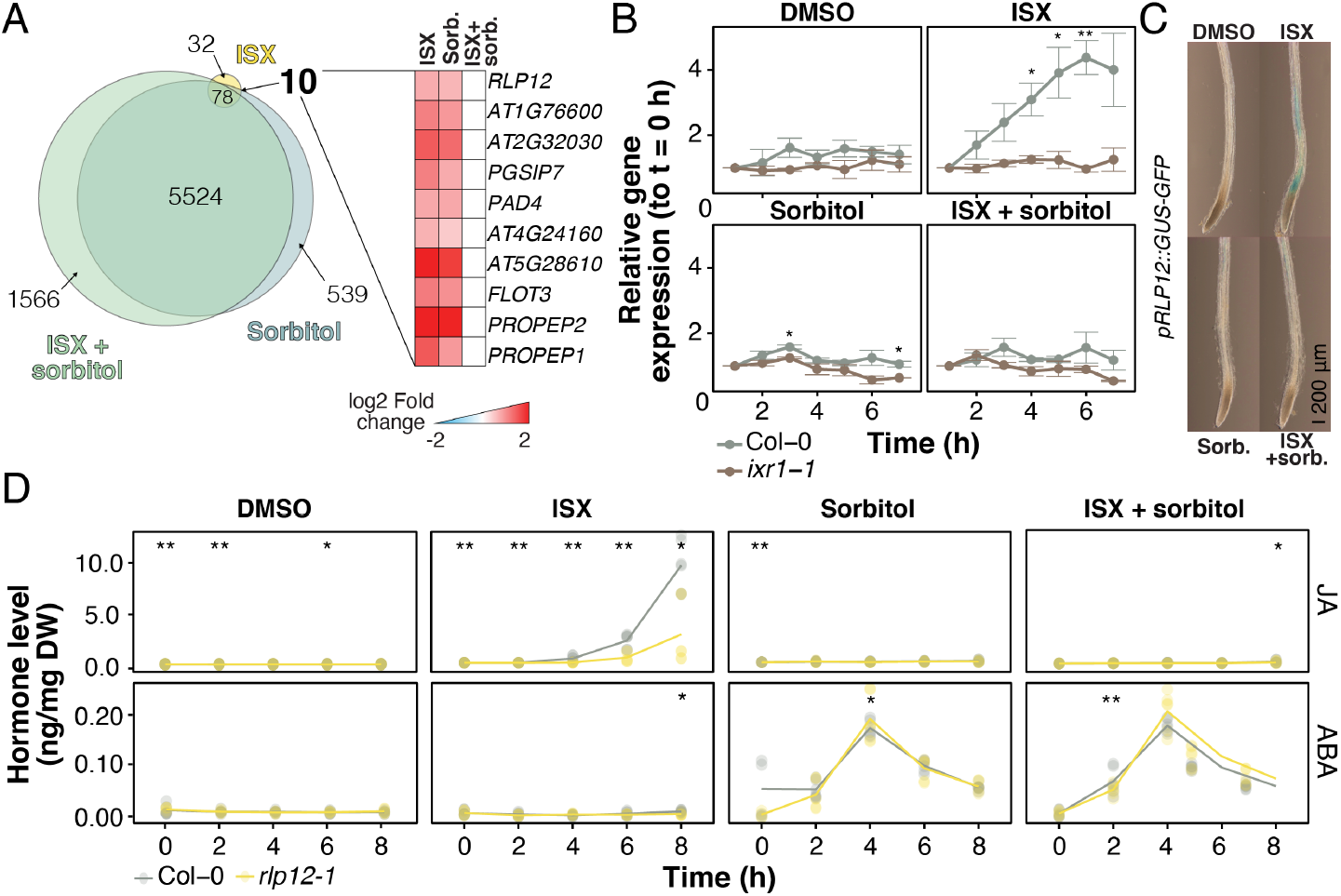
RLP12 regulates JA production induced by cell wall damage. (*A*) Results of RNAseq-based gene expression analysis of seedlings treated as indicated in figure. (*B*) Changes in expression levels of *RLP12* over time in seedlings treated as highlighted in figure. Average values ± SEM of four independent experiments with three technical replicates (*N* = 12) are shown. Asterisks indicate statistically significant differences (* *P* < 0.05, ** *P* < 0.01; ANOVA with Tukey correction for multiple comparisons). (*C*) Arabidopsis seedling roots transformed with *pRLP12::GFP-GUS* and treated as indicated in figure. (*D*) Quantification of JA and ABA levels in wildtype and *rlp12-1* seedlings. Points represent individual measurements, and lines show average values (*N* = 3). Asterisks indicate statistically significant differences (* *P* < 0.05, ** *P* < 0.01, *** *P* < 0.001, Kruskal–Wallis test).

### Cell wall integrity and ABA homeostasis modulate turgor loss points and cell wall metabolism in adult plants

The available data implicate *RLP12* and *THE1* in osmo-sensitive CWI maintenance- and possibly ABA-mediated processes in seedlings. To clarify their respective roles in more detail, we initiated a phenotypic characterisation of *RLP12* and *THE1* in seedlings and adult plants. *aba2* and *pyrobactin resistance1 like 9* (*pyr pyl*) derived plant materials were included as controls where appropriate since they affect ABA biosynthesis and signalling (41, 42). The role of *RLP12* and *THE1* in seedling root growth and ABA perception was assessed by transferring 6 days old Col-0, *pyr pyl*, *the1-1*, *the1-4* and *rlp12-1* seedlings onto plates with or without ABA (SI Appendix Fig. S4A). Seedlings from all genotypes examined exhibited root growth rates similar to Col-0. While *pyr pyl* seedling root growth was resistant to ABA, Col-0 root growth was inhibited. Root growth inhibition by ABA was more pronounced in *rlp12-1* than in Col-0. However, *the1-1* and *1-4* root growth seemed even more sensitive to ABA than *rlp12-1*. These results indicate, that THE1 and RLP12 are not essential for root growth and both *THE1* and *RLP12* seem to act as negative regulators of ABA-mediated root growth inhibition. Changes in conductance induced by ABA were characterized in Col-0, *the1-1*, *the1-4*, *rlp12-1* stomata while induction by light was also investigated in *aba2* (**Fig. 4A, B**). While pronounced differences to Col-0 were detected for *aba2* in response to light induction, none were detectable for *the1* and *rlp12-1*. However, *the1-1* and *the1-4* stomata did exhibit responses qualitatively opposite to each other upon exposure to both light and ABA. Turgor loss point has been suggested as useful tool to assess drought stress resistance and turgor levels in plants (43). We determined turgor loss points in leaves of Col-0, *aba2*, *the1-1*, *the1-4* and *rlp12-1* adult plants (**Fig. 4C**) and found that only *the1-1* leaves exhibited a pronounced increase, implying enhanced susceptibility to drought stress. Cell wall monosaccharides and cellulose were quantified in Col-0, *aba2*, *the1-1*, *the1-4* and *rlp12-1* leaves and seedlings to determine if these genes have an essential role in regulation of cell wall composition (**Fig. 4D**, SI Appendix Fig. S4C, D). No pronounced changes were detected in *the1-1, the-4,* or *rlp12-1* seedlings or leaves. In *aba2* leaves monosaccharide composition was similar to Col-0 but cellulose amounts were reduced. A similar reduction in cellulose has been reported for *aba1* plants (44). These results suggest *THE1* and *RLP12* are required for ABA-mediated processes during seedling root growth while they are not essential in adult plants.

**Fig. 4:**
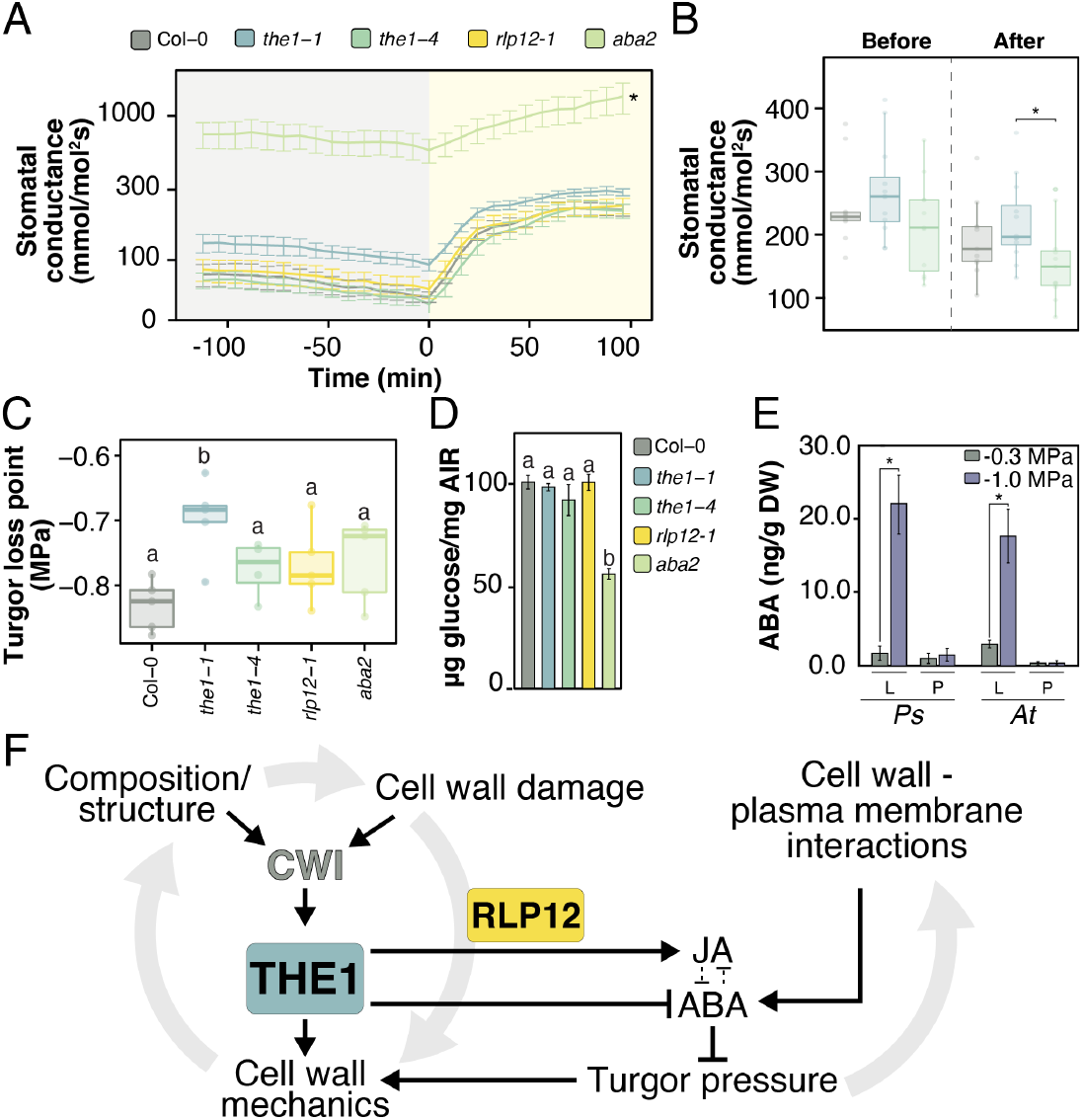
Cell wall integrity and ABA homeostasis modulate turgor loss points and cell wall metabolism in adult plants. (*A-B*) Changes in stomatal conductance (dark to light) (*A*) and in response to ABA treatment (*B*), genotypes examined indicated by colour-coding in figure (*N* = 5). Asterisks indicate statistically significant differences (* *P* < 0.05, ANOVA test with Tukey correction for multiple comparisons). (*C*) Turgor pressure loss point measurements (*N* = 5). Asterisks indicate statistically significant differences (* *P* < 0.05; ANOVA test with Tukey correction for multiple comparisons). (*D*) Cellulose content of cell walls in leaves, genotypes indicated in figure and expressed in µg of glucose per mg of alcohol insoluble fraction (AIR). (*E*) ABA levels in mock or osmoticum-treated *Pisum sativum* (*Ps*) and *Arabidopsis thaliana* (*At*) leaves (L) and protoplasts (P). Average values (*N* = 5) ± SEM. Asterisks indicate statistically significant differences (* *P* < 0.05, Student’s t-test). (*F*) Simplified model summarising the roles of THE1 and RLP12 in CWI maintenance, regulation of ABA and JA biosynthesis, as well as modulation of turgor pressure.

Based on the results presented interactions between cell wall and plasma membrane could be responsible for induction of ABA production in plants in general. To test this hypothesis Arabidopsis and *Pisum sativum* leaves and protoplasts were mock- or Polyethyleneglycol-treated and ABA production subsequently quantified (**Fig. 4E**). In mock-treated leaves and protoplasts ABA production was not enhanced. Polyethyleneglycol treatment induced ABA production in leaves from both species. However, in protoplasts from both plant species no increased ABA production was detectable. These results indicate that an intact cell wall is required for induction of ABA production in plants.

## Discussion

Here we have investigated the effects of CWD and turgor manipulation on stiffness in the Arabidopsis seedling root, determined the roles of THE1 in regulation of stiffness, ABA production and ABA mediated processes, identified RLP12 as new CWI maintenance component, and established that an intact cell wall is essential for hyper-osmotic stress-induced ABA production in plants. The results from the protoplast experiments are exciting because they provide key information enabling us to investigate the mechanism responsible for induction of ABA production, a process regulating plant adaptation to drought (21).

The model shown in **Fig. 4F** uses the knowledge generated here to outline a mechanism capable of coordinating changes in turgor pressure and cell wall stiffness in Arabidopsis. Cell wall composition, structure, CWD and stiffness are inter-connected and influence CWI. We found that stiffness in the stele of seedling roots is reduced in vivo in response to CWD (cellulose biosynthesis inhibition) and reduced turgor pressure (sorbitol). THE1 regulates stiffness changes during seedling root growth and in response to ISX while affecting those induced by ISX / sorbitol co-treatments. We did not detect global changes in wall monosaccharide composition in *the1* seedlings, suggesting that the effects on stiffness may be caused either by changes in cell wall structure or compositional changes not detectable with the analytical methods used here. THE1 acts as a positive regulator for JA in response to CWD and a negative one for ABA upon hyperosmotic stress. In *the1-1* leaves turgor loss point was reduced, suggestive of reduced turgor, which could be explained by increased ABA biosynthesis reducing turgor pressure (25). This implies that THE1 either modulates turgor pressure indirectly by modifying ABA production or THE1 could act as positive regulator of ABA production in adult plants in response to hyperosmotic (drought) stress. The results from the experiments with Arabidopsis and *P. sativum* indicate that interactions between an intact cell wall and the plasma membrane are essential for ABA production. We propose these interactions constitute an on-off switch for activation of ABA biosynthesis. Conversely, the state of the plasma membrane - cell wall continuum would finetune phytohormone biosynthesis via THE1. CWD (or hypoosmotic stress) would cause expansion and induce JA production via THE1, making THE1 a positive regulator of JA production. By contrast, plasma membrane displacement versus the wall upon hyper-osmotic shock would remove THE1-mediated repression of ABA production.

In our model, THE1 acts as positive or negative regulator dependent on changes in the cell wall - plasma membrane continuum (24). Such context-dependent behaviour has been described for FER and explained by FER acting as scaffold interacting with a large number of partners, bringing about specific activities (24). However, manipulating FER activity has profound phenotypic effects while loss of THE1 activity has only limited consequences. More importantly, FER acts as negative regulator for salt stress-induced ABA and ISX-induced JA, SA and lignin production (12, 45). This suggests that some form of redundancy compensates for loss of THE1 and that THE1 has more specialized and different functions than FER, even if both FER and THE1 contribute to stress-induced, cell wall dependent processes. Redundancy could be mediated by other molecular components involved in mechano-perception or coordination between the cell wall and cellular components (3, 4, 23, 46).

Reduced JA induction in *rlp12* seedlings after ISX treatment and ISX-induced, osmo-sensitive *RLP12* expression in the same region of the root like *THE1* indicate that *RLP12* is required for CWI maintenance. *RLP12* has been implicated previously in osmo-sensitive and biotic stress-related processes (40). The reduced JA production in *rlp12* seedlings is similar to reductions observed for *LEUCINE-RICH REPEAT KINASE FAMILY PROTEIN INDUCED BY SALT STRESS* (*LRR-KISS*) mutants (47). In contrast, mutations in Leucine rich repeat kinases, required for pattern-triggered immunity, enhance ISX-induced JA production (12, 47). A possible explanation is that *RLP12* has a specific function in CWI maintenance with effects on biotic stress response being only secondary consequences. The limited mutant phenotypes observed in *the1* and *rlp12* adult plants suggest that both genes are either primarily required at the seedling stage or redundancy may exist.

To summarize, here we have used Brillouin microscopy to investigate processes modifying cell wall stiffness in plants in vivo. The results illustrate the application potential for Brillouin microscopy and how it can complement other recently developed methods (6, 48–50). Based on the data generated, we suggest that THE1 is a key regulatory element enabling a plant cell to differentially activate phytohormone production upon exposure to either CWD or hyper-osmotic stress-induced membrane-displacement. The ability to differentially activate phytohormones in a tightly controlled manner is in turn a key pre-requisite for successful plant adaptation to a changing environment, where extreme biotic and abiotic stress situations become the norm not the exception (8, 21).

## Materials and Methods

### Plant material

*Arabidopsis thaliana* and *Pisum sativum* genotypes used in this study were obtained from the labs previously publishing them or ordered from the Nottingham Arabidopsis Stock Centre (http://arabidopsis.info/). Detailed information is listed in Extended Data Table 1.

### Plant growth conditions

Arabidopsis seedlings were grown in sterile conditions in liquid culture or on plates as described(10) with minor modifications. 30 mg of seeds were sterilized by sequential incubation with 70 % ethanol and 50 % bleach on a rotating mixer for 10 min each and washed 3 times with sterile water. Seeds were then transferred into 250 ml Erlenmeyer flasks containing 125 ml half-strength Murashige and Skoog growth medium (2.1 g/L Murashige and Skoog Basal Medium, 0.5 g/L MES salt and 1 % sucrose at pH 5.7). Seedlings were grown in long-day conditions (16 h light, 22°C / 8 h dark, 18°C) at 150 μmol m^-2^ s^-1^ photon flux density on a IKA KS501 flask shaker at a constant speed of 130 rotations per minute.

For histochemical GUS staining and microscopy imaging, Arabidopsis seeds were sterilized, and 3-5 individual seeds were placed in a well of a clear 6-well tissue-culture plate and grown in the described liquid media and growth conditions.

For experiments with protoplasts and turgor loss point measurements Arabidopsis seeds were sown directly in 10 cm pots containing a 90:10 mix of peat moss:fine grade vermiculite (v:v). Pots were sat in trays of water and covered in clear plastic wrap for 5 days until seeds had germinated, after which plastic wrap was removed and plants were thinned to one per pot. Pots were watered by immersion every three days and received a weekly watering of liquid fertilizer (Miracle-Gro, Scotts Miracle-Gro, Marysville, OH, USA). Short-day growth conditions were maintained, under a 10 h photoperiod, provided by LED lighting resulting in a PAR of 50 µmol m^-2^ s^-1^ at pot height. Temperature was maintained at 23°C. Experiments were conducted on the newest fully expanded leaves of plants that were at least 5 weeks of age.

Seeds of *P. sativum* genotype Argenteum were grown in 14 cm pots containing a 1:1 (v:v) mix of vermiculite and dolerite chips topped with 4 cm of fine, pine bark potting mix. Plants were grown in a greenhouse under a 16 h photoperiod, supplemented and extended by sodium vapor lamps which provided a minimum PAR of 300 µmol m^-2^ s^-1^ at pot height, and a 23°C/16°C day/night temperature cycle.

To measure ABA levels in protoplasts, protoplasts were prepared from approximately 20 newest fully expanded leaves of Arabidopsis genotypes and *P. sativum* as described in (51). Once isolated, protoplasts were suspended in an aqueous solution of polyethylene glycol (PEG) (MW 3000) prepared to a water potential of -0.3 MPa or -1 MPa according to (52) for 1 h. Leaves of Arabidopsis or *P. sativum* (with epidermis removed) were also excised under water and sliced into 2 mm strips then suspended PEG solutions set at either -0.3 MPa or -1 MPa for 1 h. After exposure to PEG solutions leaf samples and solutions containing protoplasts were immersed in cold -20°C 80% methanol in water (v:v) with added butylated hydroxytoluene (250 mg/L) for ABA analysis. Leaf samples were removed from PEG solutions and dried briskly on paper towel before covering in methanol solution. At least three times the volume of methanol was added to protoplast solutions. All experiments were repeated three times.

To test the effect of a brief exposure of leaf tissue to a mild reduction in leaf turgor by PEG solutions, leaves of Arabidopsis genotypes were excised under water and finely sliced into 2 mm strips. Leaf strips were then transferred to a solution of PEG set a -0.6 MPa for 20 mins. Based on pressure-volume curves this water potential should be sufficient to induce turgor loss in mesophyll cells. Following exposure to the PEG solution leaf tissue was briskly dried on paper towel before immersing in cold methanol for ABA analysis.

Plants for gas-exchange measurements were grown in 2:1 (v:v) peat:vermiculite mixture as previously described (53). Plants were grown in growth chambers (AR-66LX, Percival Scientific, USA and Snijders Scientific, Belgium) with 12 h photoperiod, 23/18 °C day/night temperature, 150 μmol m-2 s-1 light and 70% relative humidity. Plants were 24-28 days old during gas-exchange experiments.

CWD was generated in Arabidopsis seedlings using the cellulose biosynthesis inhibitor ISX at a concentration of 600nM in half-strength Murashige and Skoog growth medium. To generate hyperosmotic stress the growth medium was supplemented with 300mM sorbitol. Additional treatments include the combination of ISX and sorbitol (600nM ISX + 300mM sorbitol) and treatment with DMSO as mock control (solvent for ISX). Standard treatment duration was 6 hours unless specified differently.

### Root growth assays

Seedlings were grown for six days on plates containing growth medium with the same composition as the liquid medium but containing in addition 0.8% Agarose (Sigma). Seedlings were transferred from regular plates to plates either 10 µM ABA or no ABA (GM). Positions of root tips were marked after transfer and then roots were allowed to grow for 24 hours. After 24 hours images were taken using a Zeis Axio Zoom.V16 equipped with a PlanNeoFluar Z 1x/0.25 FWD 56mm objective lens, a 25x/10foc eyepiece and a Zeiss Axiocam506 colour. Root lengths were determined using the freehand drawing tool in Fiji Version 2.1.0. Three independent experiments were performed and for each treatment / genotype combination >15 seedling roots were measured per experiment. For calculation of relative root length, the following formula was applied:

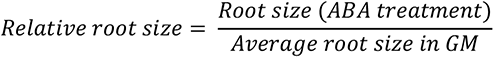

### Microscopy

#### Brillouin microscopy

Brillouin spectra were acquired using two different interferometers: time-resolved data was recorded using a VIPA-based system that offers typically 2 s acquisition times, other measurement that do not require fast scanning rates were obtained using a Tandem Fabry-Pérot interferometer. The spectrometers are both coupled with an inverted life science microscope (Eclipse Ti-U, Nikon) equipped with a micro-positioning stage. The collimated linearly polarized laser light (532-nm single mode laser, Spectra-Physics Excelsior-532), spectral linewidth < 0.01 pm) is focused onto the sample using a 20× objective lens (NA 0.35). The focal plans were adjusted on Col-0 roots, near the equatorial plan of the roots. The typical power used at the focus was 8 mW. The back-scattered light is collected by the same objective lens and is collected outside the microscope using a telescope.

The tandem Fabry-Pérot interferometer (JRS Scientific Instruments) offers a 15 GHz free spectral range sampled with a 30 MHz resolution. It is equipped with two sets of mirrors with a 95% reflectivity, and an avalanche photodiode (Count R Blue, Laser Components) with a dark count lower than 10 counts/sec (typically 2counts/sec) and a quantum efficiency of 65% at this wavelength. The magnification of the telescope that collects the backscattered light is calculated to be in accordance with the size of the 300 μm entrance pinhole of the spectrometer. A particular attention has been paid to respect the f-number (f/18) of the spectrometer to maximize photon collection on the photodiode. Each spectrum is acquired by averaging over ∼150 counts to obtain a 13 dB noise level typically (54).

The VIPA spectrometer (Light Machinery, Hyperfine HF-8999-532) is based on a 3.37 mm thick VIPA etalon (30 GHz, 500 nm to 600 nm), and is equipped with two double passed air spaced etalons to increase the contrast to about 120 dB. The fibre is directly connected to the built-in FC/PC connector of the VIPA spectrometer. The free spectral range (FSR) of the VIPA used in our work is 30 GHz, corresponding to a sampling interval of 0.11 GHz. Typical acquisitions times were 5s per point (30).

To determine the part of the Arabidopsis root responsive to CWD, 6-days-old seedlings (Col-0 and *ixr1-1* as control) grown as described above were treated for six hours with 600 nM ISX. Five regions on the roots were visually identified in each genotype for both ISX- and mock (DMSO)-treated seedlings. The regions correspond to different developmental stages (differentiation zone, elongation zone, transition zone, basal meristem and apical meristem, Extended Data Fig.1). Brillouin spectroscopy measurements were acquired radially in each zone (15 µm between points). The results were analysed, and the end of the basal meristem was chosen as target region for the next experiments (Extended Data Fig.1). To determine the effects of CWD and turgor pressure on cell wall stiffness, different treatments were performed for each genotype using the same plant material as described previously: *i*) mock treatment, *ii*) CWD (600 nM ISX); *iii*) hyperosmotic pressure (300 mM sorbitol); *iv*) combined treatment (600 nM ISX + 300 mM sorbitol). Samples (at least three per replicate, three replicates) were collected at times 0 and 6 after start of treatments for each genotype and performed as described above.

Initial data analysis was performed in Matlab (Mathworks, DE) using custom written scripts. Before the analysis, we verified that the average noise (measured in the spectral regions without any Brillouin peaks) is zero. Assuming a moderate attenuation in our systems, we fit the spectra with a Lorentzian function to determine the Brillouin frequency shift. This procedure allows reaching a 6 MHz accuracy at a 10 dB noise level. Further processing included the calculation of Brillouin Elastic Contrast, as defined in (55):

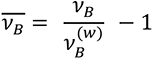

ν_B_: Measured Brillouin frequency shift

ν_B_^(w)^: Brillouin frequency shift of distilled water

This is a dimensionless parameter that normalizes the obtained Brillouin frequency shifts to those obtained for distilled water using the same setup as for the sample. Regression curves were fitted for each graph using smooth conditional medias (from ggplot2 R package) calculated using local polynomial regression fitting (LOESS) method.

#### Confocal microscopy

Fluorescence confocal laser scanning microscopy was performed using a Leica TCS SP8 (Leica Microsystems) for reporter lines *pJAZ10::YFP,* and *pRAB18::GFP-GUS*, and LSM 800 (Carl Zeiss) for reporter line *pTHE::YFP*. Seedlings were stained in the respective treatment media supplemented with 0.1% Calcofluor White Stain solution for 30 min and rinsed for 5-10 times in pure treatment media before imaging. Seedlings were mounted in treatment media. Image acquisition was done with HC PL APO CS 10x/0.40 NA dry and HC PL APO CS2 20x/0.75 NA dry objectives in Leica TCS SP8 (Leica Microsystems). Calcofluor White was excited at 405 nm laser and emission was detected between 425 – 475 nm. YFP/GFP was excited at 514 nm and emission was detected between 519 – 553 nm in Leica TCS SP8 (Leica Microsystems). The acquisition for LSM 800 (Carl Zeiss) was done with Plant-Neofluar 10x/0.3 NA dry objective in LSM 800 (Carl Zeiss), and samples were imaged using the excitation laser line 405 nm and emission was detected 400-470 nm for Calcofluor White fluorescence detection, and the excitation laser line 488 nm and emission spectrum was detected 507-700 nm for YFP. Images obtained from Leica TCS SP8 (Leica Microsystems) were processed in Fiji (ImageJ), and images obtained from LSM 800 (Carl Zeiss) were processed as single plane maximum intensity projections of 3 optical sections in Zen Blue 3.3 software (Carl Zeiss).

### Histochemical GUS staining

7-day old seedlings were mock-, ISX-, sorbitol- or an ISX/Sorbitol-treated for 6h and submerged into GUS staining solution (50 mM phosphate buffer (pH = 7), 2 mM ferricyanide K_3_Fe(CN)_6_, 2 mM ferrocyanide (K_4_Fe(CN)_6_*3H_2_O), 2 mM 5-Bromo-4-chloro-3-indolyl β-D-glucopyranoside (X-Glc), 0.2% Triton X-100) immediately after. The samples were incubated for 60 min at 37°C in darkness and transferred to 70% ethanol for 2 days (at RT) prior to imaging. Seedlings were imaged with a Zeiss Axio Zoom.V16 equipped with a PlanNeoFluar Z 1x/0.25 FWD 56mm objective lens, a 25x/10foc eyepiece and a Zeiss Axiocam506 colour. For image acquisition, intensity of the Zeiss CL 9000 LED light source was set to 21%. Other settings used were a fully opened aperture, motorized zoom factor of 5.1x (resulting in a final root magnification of 128x), and an exposure time of 150 ms.

### Phytohormone measurements

JA and ABA were analysed as described(56), with minor modifications. Arabidopsis seedlings were flash-frozen in liquid nitrogen and freeze-dried for 24 h. 6-7 mg aliquots of freeze-dried seedlings were ground with 5 mm stainless steel beads in a Qiagen Tissue Lyser II for 2 min at 25 Hz. Shaking was repeated after addition of 400 μL extraction buffer (10 % methanol, 1 % acetic acid) with internal deuterated standards (50 ng/ml Jasmonic-d_5_ acid, 50 ng/ml Salicylic-D_4_ acid and 10 ng/ml Abscisic-D_6_ acid); CDN Isotopes, Pointe-Claire, Canada) before samples were incubated on ice for 30 min and centrifuged for 10 min at 16,000 g and 4°C. Supernatants were transferred into fresh tubes and centrifuged 3 times to remove all debris prior to LC-MS/MS analysis. An extraction control not containing plant material was treated equally to the plant samples.

Chromatographic separation was carried out on a Waters ACQUITY I Class UPLC system, equipped with a Waters Cortecs C18 column (2.7 mm, 2.1 x 100 mm). Water was used as mobile phase A and acetonitrile was used as mobile phase B, both containing 0.1 % formic acid. The linear mobile phase gradient was adapted to a total run time of 7 min: 0-4 min 20 % to 95 % B, 4-5 min 95 % B, 5-7 min 95 % to 20 % B; flow rate 0.4 ml/min. The column temperature was maintained at 35 °C, the injection volume was 5 μL.

For hormone detection and quantification, a Waters Xevo TQ-XS triple quadrupole mass spectrometer was employed. The mass spectrometer was equipped with an ESI source operated in negative mode and MS conditions were set as follows: capillary voltage -1.8 kV; cone voltage 25 V; source offset voltage 30 V; source temperature 150 °C; desolvation temperature 500 °C; cone gas flow 150 L/h; desolvation gas flow 1000 L/ h; collision gas flow 0.15 ml/min; nebulizer gas flow 6 Bar. The spectra were acquired in multiple reactions monitoring mode and following precursor-product ion transitions were used for quantification: JA 209.10 > 58.98, D_5_-JA 214.10 > 62.00, SA 136.97> 93.20, D_4_-SA 140.97 > 97.20, ABA 263.00 > 153.00, D_6_-ABA 269.00 > 159.00.

Individual stock solutions of phytohormones were prepared in methanol. A series of standard mixes in the range of 0.2–1000 ng/ml, prepared in extraction solvent containing deuterated internal standards, were used to build up calibration curves.

ABA levels in Arabidopsis and *P. sativum* protoplasts and leaves were quantified physico-chemically with an added internal standard according to the method previously described (57) using an Agilent 6400 Series Triple Quadrupole liquid chromatograph tandem mass spectrometer. To remove excess PEG from samples, the sample in 200 µL of 2% acetic acid in water (v:v) was purified by partitioning twice with 100 µL of diethyl ether, and with the organic phase containing the ABA dried to completeness at 40°C, after the sample was resuspended in 200 µl of 2% acetic acid in water (v:v). ABA samples in *P. sativum* were prepared similarly to those of *Arabidopsis* but were quantified physico-chemically with an added internal standard according using a UPLC-MS/MS as previously described(58).

### Generation of transgenic lines

Genomic DNA (gDNA) was extracted from 7-day old Arabidopsis Col-0 seedlings with E.Z.N.A Plant DNA kit (Omega). The full-length *RLP12* promoter (*pRLP12*) was amplified using gDNA as template with primer pairs containing AttB1/2 recombination sites (see Extended Data Table 2). *pRLP12* was inserted through Gateway Recombination Cloning Technology (Life Technologies) in the binary pFAST-G04 (containing GFP-GUS fusion) vector (59) used as destination. Upon Agrobacterium mediated transformation, we selected, by using Zeiss Axio Zoom V16 equipped with fluorescence filters for GFP, 7 independent transgenic stable T1 plants (in Col-0 background). Experiments were carried on two homozygous T3 lines carrying a single insertion.

### Transcriptomics

Total RNA was extracted using a Spectrum Plant Total RNA Kit (Sigma-Aldrich). RNA integrity was assessed using an Agilent RNA6000 Pico Kit. Sequencing samples were prepared and analysed as described previously(13). The intersection of the DE genes between the treatments was visualized using a Venn diagram produced with R package BioVenn (60).

### qRT-PCR-based gene expression analysis

Total RNA was isolated using a Spectrum Plant Total RNA Kit (Sigma-Aldrich). 2 mg of total RNA were treated with RQ1 RNase-Free DNase (Promega) and processed with the ImProm-II Reverse Transcription System (Promega) for cDNA synthesis. qRT-PCR was performed using a LightCycler 480 SYBR Green I Master (Roche), and primers (Extended Data Table 2) diluted according to manufacturer specifications. *ACT2* is used as reference in all experiments(61). The expression levels of each gene were determined using the Pfaffl method(62). All experiments were repeated at least three times.

### Stomatal conductance measurements

Stomatal conductance of plants was measured using a rapid-response gas-exchange measurement device consisting of eight thermostated flow-through whole-rosette cuvettes. Plants were inserted into the cuvettes in the morning and when their stomatal conductance had stabilized, darkness was applied for two hours. Subsequently light was applied to register light-induced increase of stomatal conductance within next two hours. To study ABA-induced stomatal closure plants were sprayed with 5 µM ABA solution with 0.012% Silwet L-77 (Duchefa) and 0.05% ethanol as described previously(63). ABA-induced changes in stomatal conductance were examined at gs18 and gs0, where gs0 is the pre-treatment stomatal conductance and gs18 is the value of stomatal conductance 18 min after ABA spraying.

### Pressure-volume curve analysis

Pressure-volume curves were used to determined leaf turgor loss point in five leaves from each *Arabidopsis* genotype according to the methods of (64). Briefly, the petioles of leaves were excised under water and rehydrated overnight in a sealed bag. Leaves were imaged to determine leaf area, and then concurrent and periodic measurements of leaf mass (±0.0001g) and water potential were conducted using a Scholander pressure chamber with microscope to determine the balance pressure, until at least three measurements had been collected in each leaf past turgor loss point. Leaves were then dried at 70°C and weighed to determine dry mass.

### Cell wall analysis

Arabidopsis seedlings or leaves were lyophilized and ball-milled in a Retsch mixer mill. All samples were extracted three times with 70 % ethanol at 70°C, washed with acetone and dried in a vacuum concentrator. The alcohol insoluble residue (AIR) was weighed out in 2 ml screw caps tubes and used for extraction of neutral cell wall sugars, uronic acids and cellulose as described (65). High-performance anion-exchange chromatography with pulsed amperometric detection (HPAEC-PAD) was performed on a biocompatible Knauer Azura HPLC system, equipped with an Antec Decade Elite SenCell detector. Monosaccharides were separated on a Thermo Fisher Dionex CarboPac PA20 column with a solvent gradient of (A) water, (B) 10mM NaOH and (C) 700mM NaOH at 0.4 ml/min flow rate and 40°C column/detector temperature. 0 to 25 min: 20% B, 25 to 28 min: 20 to 0% B, 0 to 70% C, 28 to 33 min: 70 % C, 33 to 35 min: 70 to 100% C, 35 to 38 min: 100% C, 38 to 42 min: 0 to 20% B, 100 to 0% C, 42 to 60 min: 20% B.

### Statistical Analysis

Statistical significance of data under normal distribution was tested using either Student’s *t*-test or one-way ANOVA followed by post-hoc analysis with Tukey’s HSD test. Statistical details of experiments are specified in the figure legends. In the case of data not following normal distribution (as indicated by Saphiro-Wilk test at α = 0.05(66)), a non-parametric test (Wilcoxon rank sum test, multiple comparisons corrected with continuous correction) was performed. Statistically significant differences are indicated by * *P* < 0.05, ** *P* < 0.01, *** *P* < 0.001 paired comparisons and different letters are used for multiple comparisons at α = 0.05. Statistical analysis were performed in R(67) using the packages ggplot2(68), dplyr(69), multcomp(70), and lsmeans(71).

## Acknowledgments

The authors would like to thank Luis Alonso-Baez for critical reading of the manuscript. We acknowledge the use of the facilities of the Bindley Bioscience Center, a core facility of the National Institute for Health-funded Indiana Clinical and Translational Sciences Institute. We thank David Nichols for running ABA samples by UPLC-MS/MS.

---

**Fig. S1: Determination of cell wall stiffness in *Arabidopsis thaliana* roots using Brillouin light scattering microscopy.** (*A*) Diagram showing the different regions in an *A. thaliana* root. Brillouin measurements were performed in the longitudinal axis, starting 250 µm away from the root tip (blue arrow). (*B*) Brillouin elastic contrast measurements (νB) across the longitudinal axis of different *A. thaliana* wildtype (Col-0) roots before and after treatment with isoxaben. Dots represent individual measurements (*N* = 3), whereas the lines represent the corresponding regression curves (LOESS adjustment) with confidence interval (shadowed). Brillouin elastic contrast is calculated as the ratio between measured frequency shift and the Brillouin frequency shift of distilled water in the same conditions, and it is a dimensionless parameter. (*C*) Light microscopy images of wildtype seedling roots used for Brillouin microscopy-based analysis of cell wall stiffness. Different treatments are indicated in figure. Red arrows highlight the reference point for measurements with results presented in Fig. 1 (0 µm in the X-axis and 450 µm from root tip).

**Fig. S2: Hyperosmotic and THE1-based signalling modulate each-others transcript levels.** Expression levels of *RAB18* and *TCH4* in whole seedlings after 6h with treatment and genotypes as indicated in the figure. Values are normalized for each gene to the standard *ACT2*. Average values ± SEM from three independent experiments with three technical replicates per experiment (*N* = 9) are shown. Letters indicate within-group differences between genotypes (ANOVA, *P* < 0.05, Tukey correction for multiple comparisons)

**Fig. S3: Different alleles of *RECEPTOR-LIKE PROTEIN 12* (*RLP12*) exhibit similar effects on CWD-induced phytohormone production.** (*A*) Position of T-DNA insertions in RLP12. (*B*) JA and ABA levels whole seedlings, with genotypes and treatments (6 hours) indicated in figure. Letters indicate within-group differences between genotypes (ANOVA, *P* < 0.05, Tukey correction for multiple comparisons).

**Fig. S4: Results of the phenotypic characterization of different genotypes implicated in CWI maintenance and ABA metabolism.** (*A*) Root growth assays of seedlings grown on growth medium and transferred to ABA containing medium. Genotypes are indicated in figure and data are presented as relative root length compared to mock conditions (GM, *N* > 10). Asterisks indicate statistically significant differences between non-exposed (GM) and ABA-exposed seedlings (* *P* < 0.05, ** *P* < 0.01, *** *P* < 0.001; ANOVA with Tukey correction for multiple comparisons). Letters indicate significant within-group differences between genotypes exposed to the same treatment (* *P* < 0.05, ANOVA test with Tukey correction for multiple comparisons). (*B*) Analysis of cell wall composition in leaves (*N* = 4) with genotypes indicated in figure. Monosaccharide composition (in mole percent, Mol. %) of dry alcohol insoluble fraction (AIR). Ara: arabinose; Fuc: fucose. Gal: galactose; GalUA: galacturonic acid; Rha: rhamnose; Xyl: xylose). Letters indicate significant within-group differences between genotypes (* *P* < 0.05, ANOVA test with Tukey correction for multiple comparisons). (*C*) Analysis of cell wall composition in seedlings with genotypes indicated in figure. Monosaccharide composition (in mol percent, Mol. %) of dry alcohol insoluble fraction (AIR) and µg of glucose per mg of AIR were determined. Ara: arabinose; Fuc: fucose. Gal: galactose; GalUA: galacturonic acid; Rha: rhamnose; Xyl: xylose). Average values (*N* = 4) ± SEM. Letters indicate significant within-group differences between genotypes (* *P* < 0.05, ANOVA test with Tukey correction for multiple comparisons).

**Supplementary Data Table 1:**
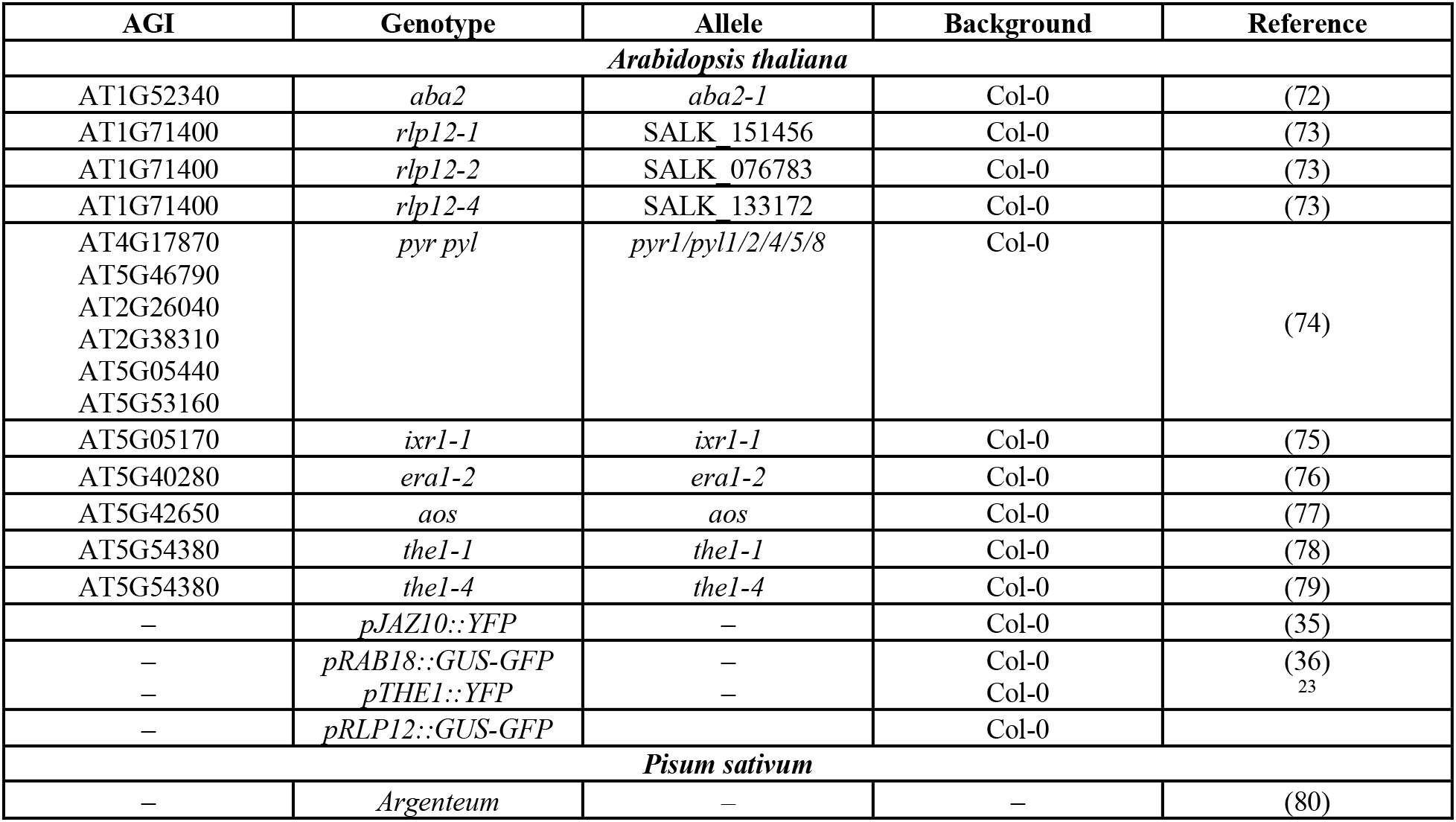
Plant material used in this study.

**Supplementary Data Table 2:**
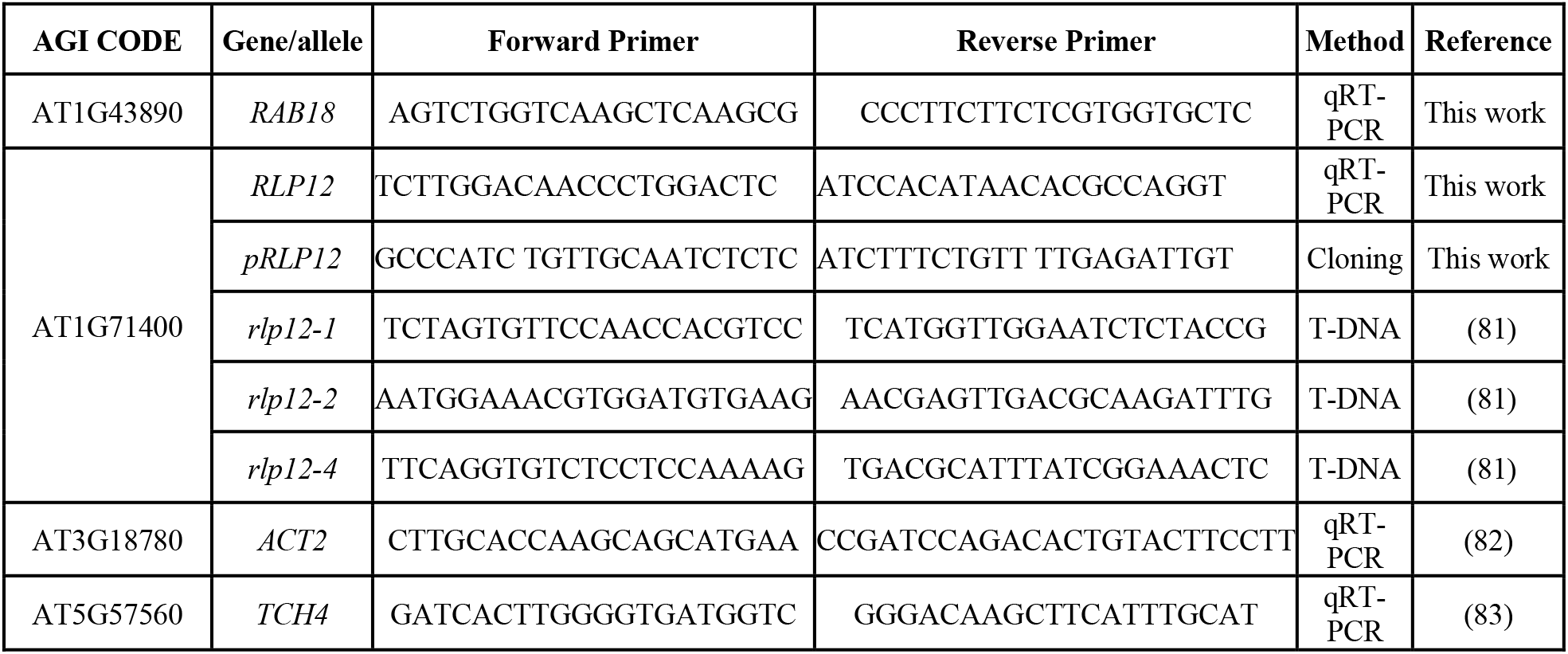
Primers used in this study.

