## Supplemental figures 1 for "THESEUS1 modulates cell wall stiffness and abscisic acid production in *Arabidopsis thaliana*"

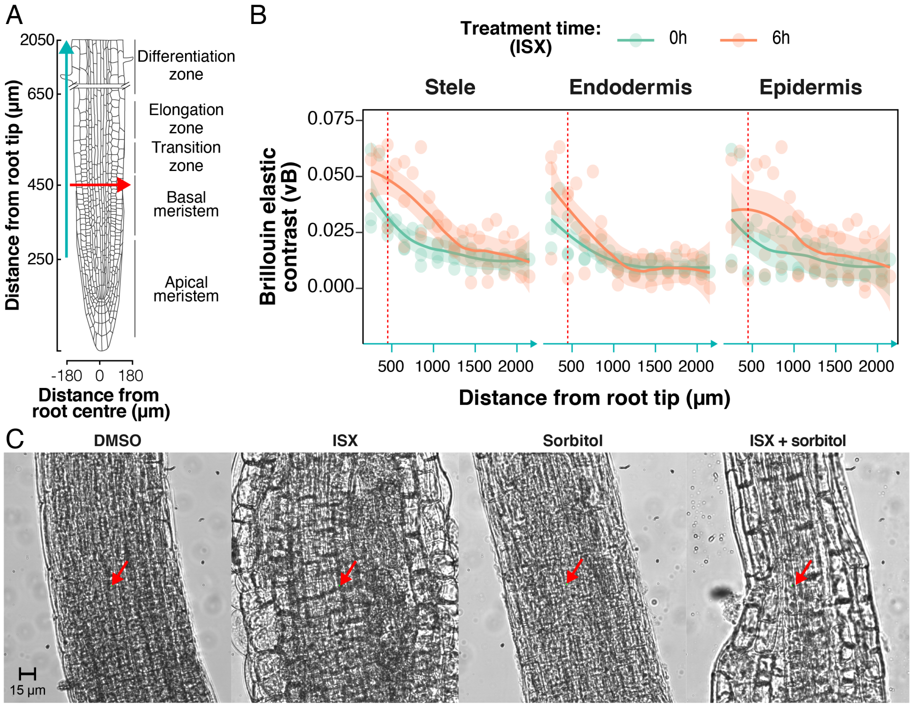


- **Fig. S1: Determination of cell wall stiffness in *Arabidopsis thaliana* roots using Brillouin light scattering microscopy.** (*A*) Diagram showing the different regions in an *A. thaliana* root. Brillouin measurements were performed in the longitudinal axis, starting 250 µm away from the root tip (blue arrow). (*B*) Brillouin elastic contrast measurements (vB) across the longitudinal axis of different *A. thaliana* wildtype (Col-0) roots before and after treatment with isoxaben. Dots represent individual measurements (*N* = 3), whereas the lines represent the corresponding regression curves (LOESS adjustment) with confidence interval (shadowed). Brillouin elastic contrast is calculated as the ratio between measured frequency shift and the Brillouin frequency shift of distilled water in the same conditions, and it is a dimensionless parameter. (*C*) Light microscopy images of wildtype seedling roots used for Brillouin microscopy-based analysis of cell wall stiffness. Different treatments are indicated in figure. Red arrows highlight the reference point for measurements with results presented in Fig. 1 (0 µm in the X-axis and 450 µm from root tip).

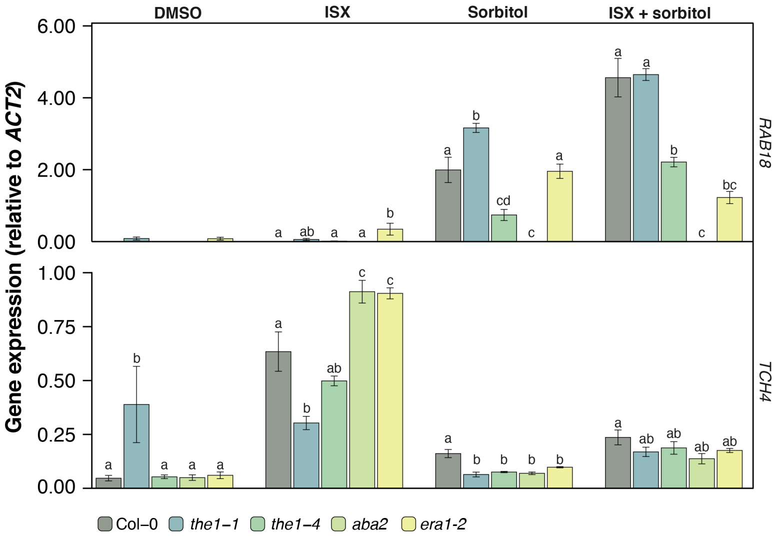


- **Fig. S2: Hyperosmotic and THE1-based signalling modulate each-others transcript levels.** Expression levels of *RAB18* and *TCH4* in whole seedlings after 6h with treatment and genotypes as indicated in the figure. Values are normalized for each gene to the standard *ACT2*. Average values ± SEM from three independent experiments with three technical replicates per experiment ($N$ = 9) are shown. Letters indicate within-group differences between genotypes (ANOVA, $P < 0.05$, Tukey correction for multiple comparisons)

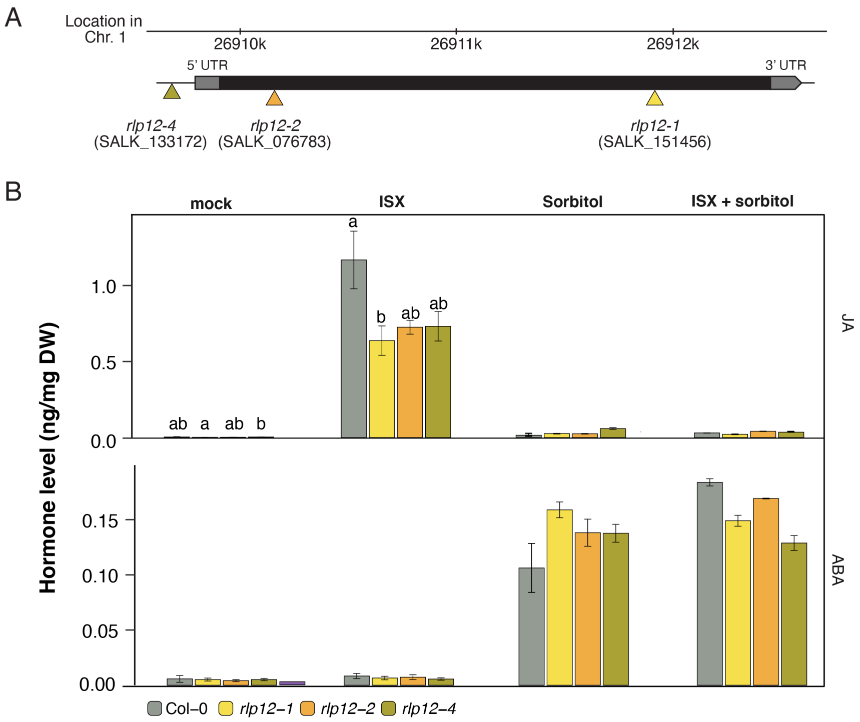


- **Fig. S3: Different alleles of *RECEPTOR-LIKE PROTEIN 12* (*RLP12*) exhibit similar effects on CWD-induced phytohormone production.** (*A*) Position of T-DNA insertions in RLP12. (*B*) JA and ABA levels whole seedlings, with genotypes and treatments (6 hours) indicated in figure. Letters indicate within-group differences between genotypes (ANOVA, $P < 0.05$, Tukey correction for multiple comparisons).

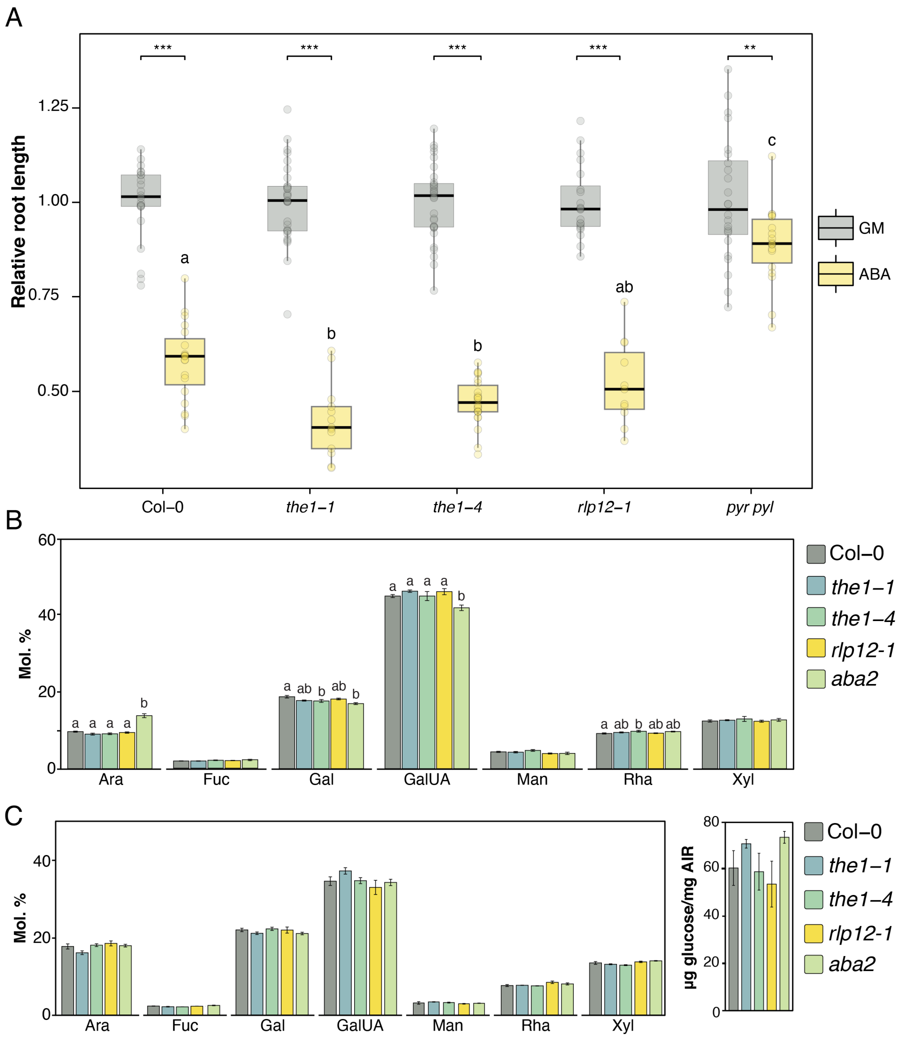


- **Fig. S4: Results of the phenotypic characterization of different genotypes implicated in CWI maintenance and ABA metabolism.** (*A*) Root growth assays of seedlings grown on growth medium and transferred to ABA containing medium. Genotypes are indicated in figure and data are presented as relative root length compared to mock conditions (GM, $N > 10$). Asterisks indicate statistically significant differences between non-exposed (GM) and ABA-exposed seedlings (* $P < 0.05$, ** $P < 0.01$, *** $P < 0.001$; ANOVA with Tukey correction for multiple comparisons). Letters indicate significant within-group differences between genotypes exposed to the same treatment (* $P < 0.05$, ANOVA test with Tukey correction for multiple comparisons). (*B*) Analysis of cell wall composition in leaves ($N = 4$) with genotypes indicated in figure. Monosaccharide composition (in mole percent, Mol. %) of dry alcohol insoluble fraction (AIR). Ara: arabinose; Fuc: fucose. Gal: galactose; GalUA: galacturonic acid; Rha: rhamnose; Xyl: xylose). Letters indicate significant within-group differences between genotypes (* $P < 0.05$, ANOVA test with Tukey correction for multiple comparisons). (*C*) Analysis of cell wall composition in seedlings with genotypes indicated in figure. Monosaccharide composition (in mol percent, Mol. %) of dry alcohol insoluble fraction (AIR) and µg of glucose per mg of AIR were determined. Ara: arabinose; Fuc: fucose. Gal: galactose; GalUA: galacturonic acid; Rha: rhamnose; Xyl: xylose). Average values ($N = 4$) ± SEM. Letters indicate significant within-group differences between genotypes (* $P < 0.05$, ANOVA test with Tukey correction for multiple comparisons).
